# Cell-free chromatin particles activate immune checkpoints in human T cells: Implications for cancer therapy

**DOI:** 10.1101/2023.06.09.544311

**Authors:** Snehal Shabrish, Kavita Pal, Naveen Kumar Khare, Dharana Satsangi, Aishwarya Pilankar, Vishalkumar Jadhav, Sushma Shinde, Nimisha Raphael, Gaurav Sriram, Relestina Lopes, Gorantla V. Raghuram, Harshali Tandel, Indraneel Mittra

## Abstract

Immune checkpoint blockade is an exciting breakthrough in cancer therapy, but how immune checkpoints are activated is unknown. We have earlier reported that cell-free chromatin particles (cfChPs) that circulate in the blood, or those that are released locally from dying cells, are readily internalized by healthy cells with biological consequences. Here we show that treatment of human lymphocytes with cfChPs isolated from sera of cancer patients led to marked activation of immune checkpoints viz. PD-1, CTLA-4, LAG-3, NKG2A, and TIM-3. Concurrently activated were stress-related markers cJun, cFos, JunB, FosB, NFКB, and EGR1. The above immune checkpoints were also activated when lymphocytes were treated with cfChPs released from dying HeLa cells; the latter could be abrogated by three cfChPs deactivating agents. These results suggest that immune checkpoints are activated by lymphocytes as stress response to cfChPs. Simultaneous downregulation of multiple immune checkpoints may herald a new approach to immunotherapy of cancer.

**Statement of Significance:** We show that cell-free chromatin particles (cfChPs) that circulate in the blood of cancer patients, or those released from dying cancer cells, simultaneously activate five immune checkpoints as a stress response by human lymphocytes. Activation of checkpoints was abrogated by cfChPs deactivating agents suggesting a novel approach to cancer treatment.

## Introduction

Immune checkpoint molecules prevent the immune system from indiscriminately attacking self-cells. Immunologically altered cancer cells protect themselves from elimination by activating immune checkpoints (1). Targeting activated immune checkpoints with specific inhibitors has revolutionized immunotherapy of cancer (2,3). Several immune checkpoint inhibitors have now been approved by FDA for the treatment of a variety of cancers (4). Although much has been reported on immune checkpoint biology and immune therapy, how immune checkpoints are activated by lymphocytes has not been elucidated.

Several hundred billion to a trillion cells die in the body everyday (5,6) and the fragmented chromosomal material in the form of cell-free chromatin particles (cfChPs) enter into the extracellular compartment of the body, including into the circulation (7–9). We have reported that cfChPs that circulate in blood of cancer patients, and those that are released locally from dying cancer cells, can readily enter into healthy cells to activate two hallmarks of cancer viz. DNA damage and inflammation (10,11). Since immune escape by the way of activation of immune checkpoints is another critical hallmark of cancer (12), we investigated whether cfChPs might be the agents that also activate immune checkpoints. We approached this question in two ways: by directly treating isolated human T-cells with cfChPs isolated from sera of cancer patients; and by using a co-culture system employing hypoxia-induced dying HeLa cells as cfChPs donors and human T-cells as recipients. We found that under both these conditions cfChPs were capable of activating five immune checkpoints as also multiple stress-related markers. Immune checkpoints against which inhibitors are in therapeutic use, viz. PD-1, CTLA-4, and LAG-3, could be markedly downregulated by concurrent treatment with three different agents that have the property of deactivating/inactivating cfChPs. These agents included anti-histone antibody complexed nanoparticles (CNPs), DNase I and a novel pro-oxidant combination of small quantities of the nutraceuticals Resveratrol and copper (R-Cu). Our results suggest that immune checkpoints are activated as a response to cfChPs-induced cellular stress. Simultaneous downregulation of multiple immune checkpoints by cfChPs deactivators heralds the prospect of a novel approach to immunotherapy of cancer. Their low cost and lack of toxicity make them attractive agents for urgent evaluation in clinical trials.

## Results

### cfChPs are readily internalized by human lymphocytes

In confirmation of our earlier report (11), treatment of human PBMCs with fluorescently dually labelled cfChPs isolated from sera of cancer patients resulted in their rapid uptake by human PBMCs by 2h **(Supplementary Fig. S1)**.

### cfChPs up-regulate immune checkpoints

Purified CD4^+^T and CD8^+^T cells were treated with cfChPs isolated from sera of cancer patients (10ng) and time course analyses were performed by qRT-PCR to detect mRNA expression of 5 immune checkpoints. All five immune checkpoints, viz. PD-1, CTLA-4, LAG-3, NKG2A and TIM-3, were found to have been markedly up-regulated, albeit at different time points **(Fig. 1A)**. These results were validated by immuno-fluorescence and flow cytometry **(Fig. 1B & C; Supplementary Fig. S2 & S3)**. This data provides evidence that cfChPs that circulate in blood of cancer patients can directly activate immune checkpoints in human CD4+ and CD8+ T cells.

**Figure 1:**
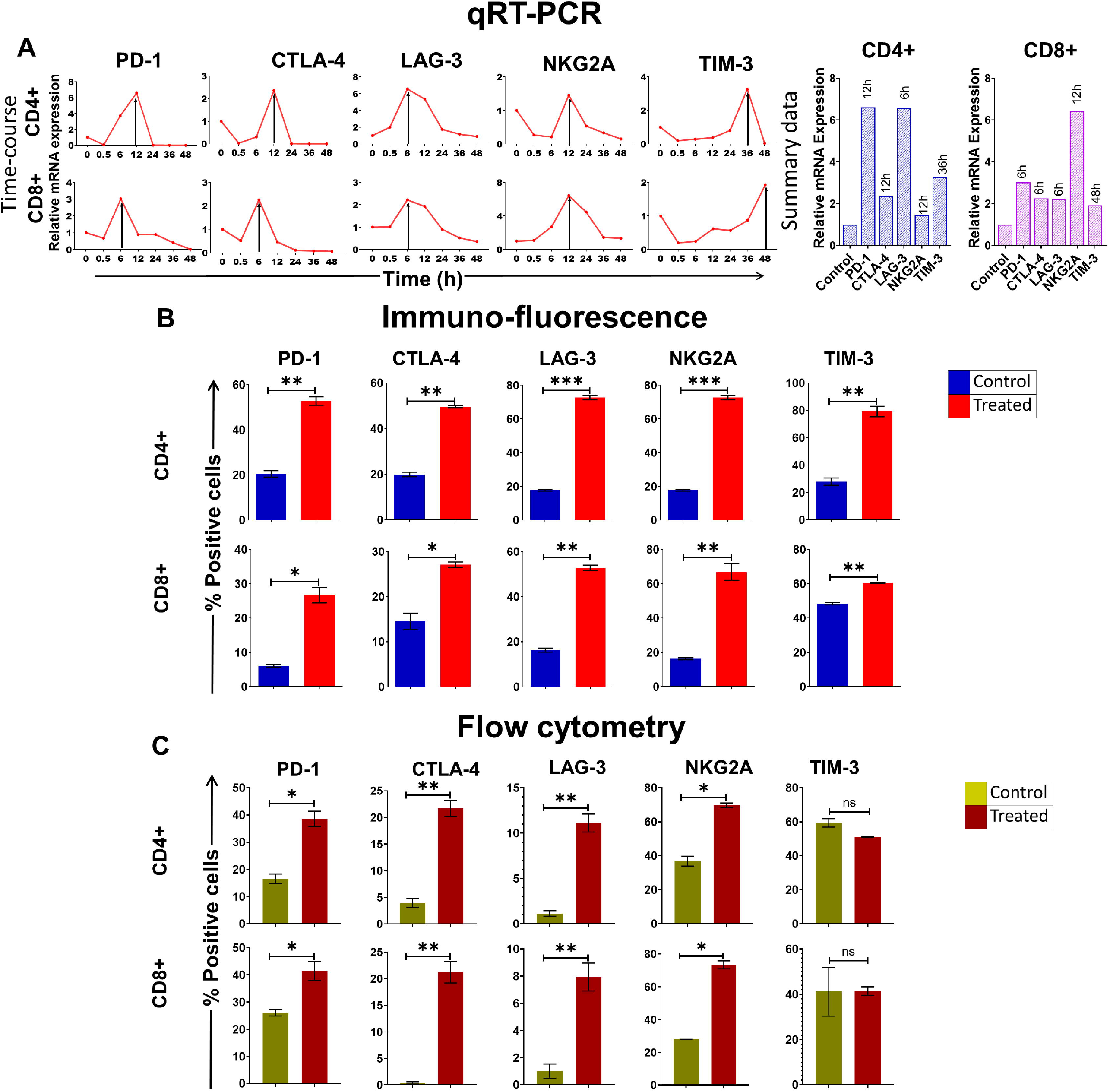
Upregulation of immune checkpoints on human lymphocytes following treatment of cfChPs isolated from sera of cancer patients. **(A)** Line graphs showing time course analysis of mRNA expression of immune checkpoints detected by qRT-PCR in purified CD4+T cells and CD8+T cells treated with cfChPs (10ng). The expression fold change was analyzed using a comparative C_T_ method [2^(-ΔΔC_T_)]. The histograms represent relative mRNA expression at respective peak time points on CD4+T cells and CD8+T cells. **(B)** Histograms depict the results of quantitative Immunofluorescence (IF) analysis of percent positive cells for immune checkpoints on CD4+T and CD8+T-cells at peak time points of mRNA expression. Five hundred cells were examined in duplicate slides and percent biomarker-positive cells were recorded. **(C)** Histograms depict the results of quantitative flow cytometry analysis of surface expression of immune checkpoint on CD4+T and CD8+T-cells. All experiments were performed in duplicates and histograms represent mean ± SEM values. Statistical analyses were performed using a two-tailed student’s unpaired t-test (GraphPad Prism 8). * p<0.05, ** p<0.01, **** p<0.0001.

### cfChPs upregulate immune checkpoints *in vivo*

In order to replicate the above findings in *in vivo* we injected cfChPs (100ng) isolated from sera of cancer patients intravenously into mice. A time course analysis by flow cytometry of isolated splenocytes detected markedly increased surface expression of four out of five immune checkpoints on CD4^+^T and CD8^+^T cells. Time points at which immune checkpoints were activated included PD-1 (6h), CTLA-4 (6h), NKG2A (6h) and LAG-3 (24h). Expression of TIM-3 was not detected during the course of the experiment lasting 96h **(Fig. 2)**.

**Figure 2:**
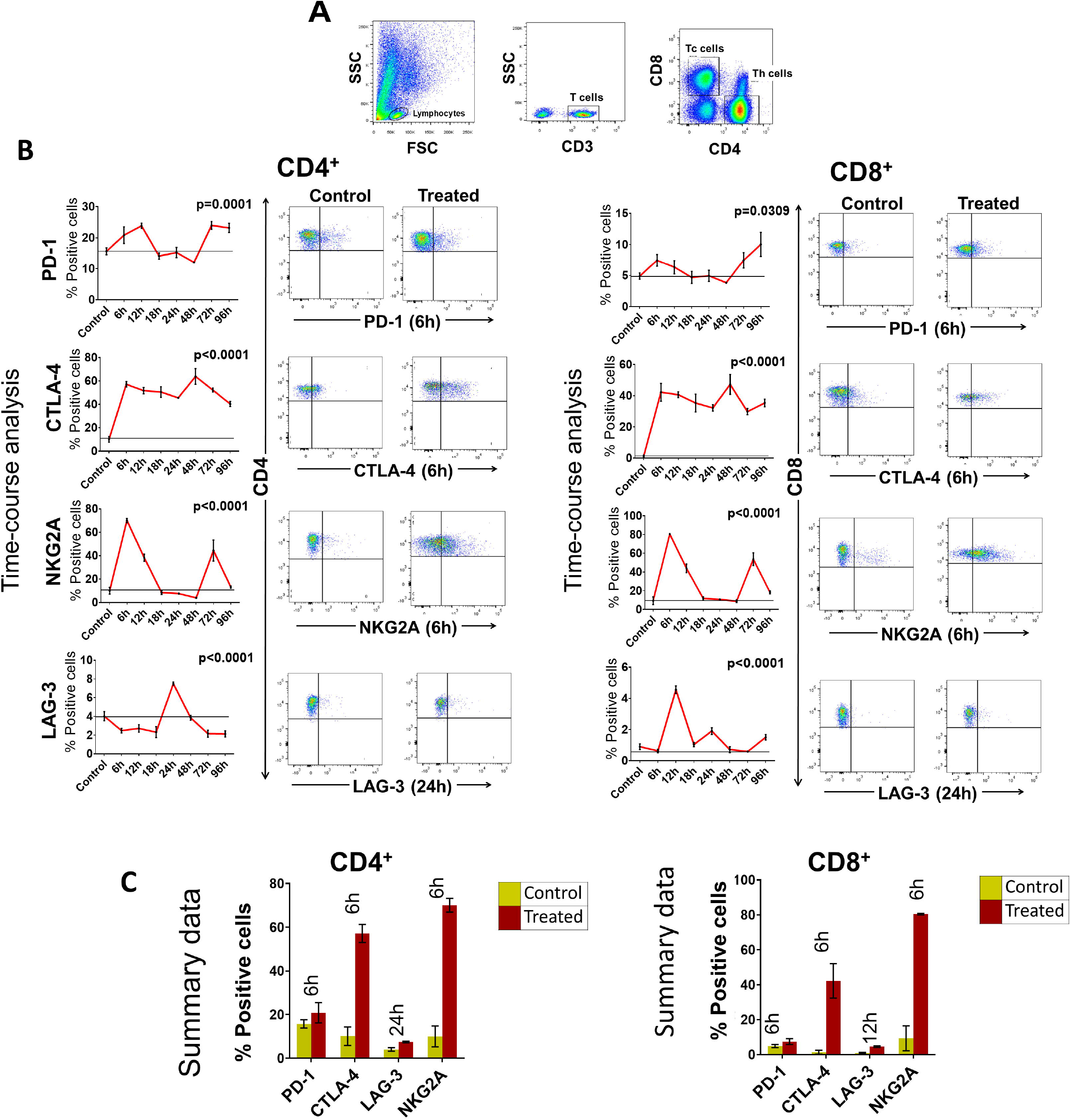
Upregulation of immune checkpoints on splenic lymphocytes of mice following intravenous injection of cfChPs (100ng) isolated from sera of cancer patients. **(A)** Gating of splenic lymphocytes on forward/side scatter, T-cells were identified by using CD3 antibodies, Th cells were identified by using CD4 antibodies and Tc cells were identified by using CD8 antibodies. **(B)** Line graphs showing results of time course analysis of surface expression of various immune checkpoints on CD4^+^T cells and CD8^+^T cells by flow cytometry (N=3 at each time point) (left-hand panels). Representative flow cytometry plots at respective peak time points are given in right-hand panels. Percent expression of each immune checkpoint was compared with untreated controls. **(C)** Histograms represent immune checkpoint expression on CD4+T cells and CD8+T cells at respective peak time points determined by flow cytometry in control and treated mouse splenocytes. Statistical analyses were performed using Bonferroni’s multiple comparisons test (GraphPad Version 8).

### cfChPs induce cellular stress

qRT-PCR analysis for detection of mRNA expression by human PBMCs following cfChPs treatment (10ng) of six stress-related markers viz. NFКB, JunB, c-Jun, c-Fos, FosB and EGR-1 was undertaken in a time-course analysis. All six stress-related markers were found to have been up-regulated, albeit at different time points **(Fig. 3)**.

**Figure 3:**
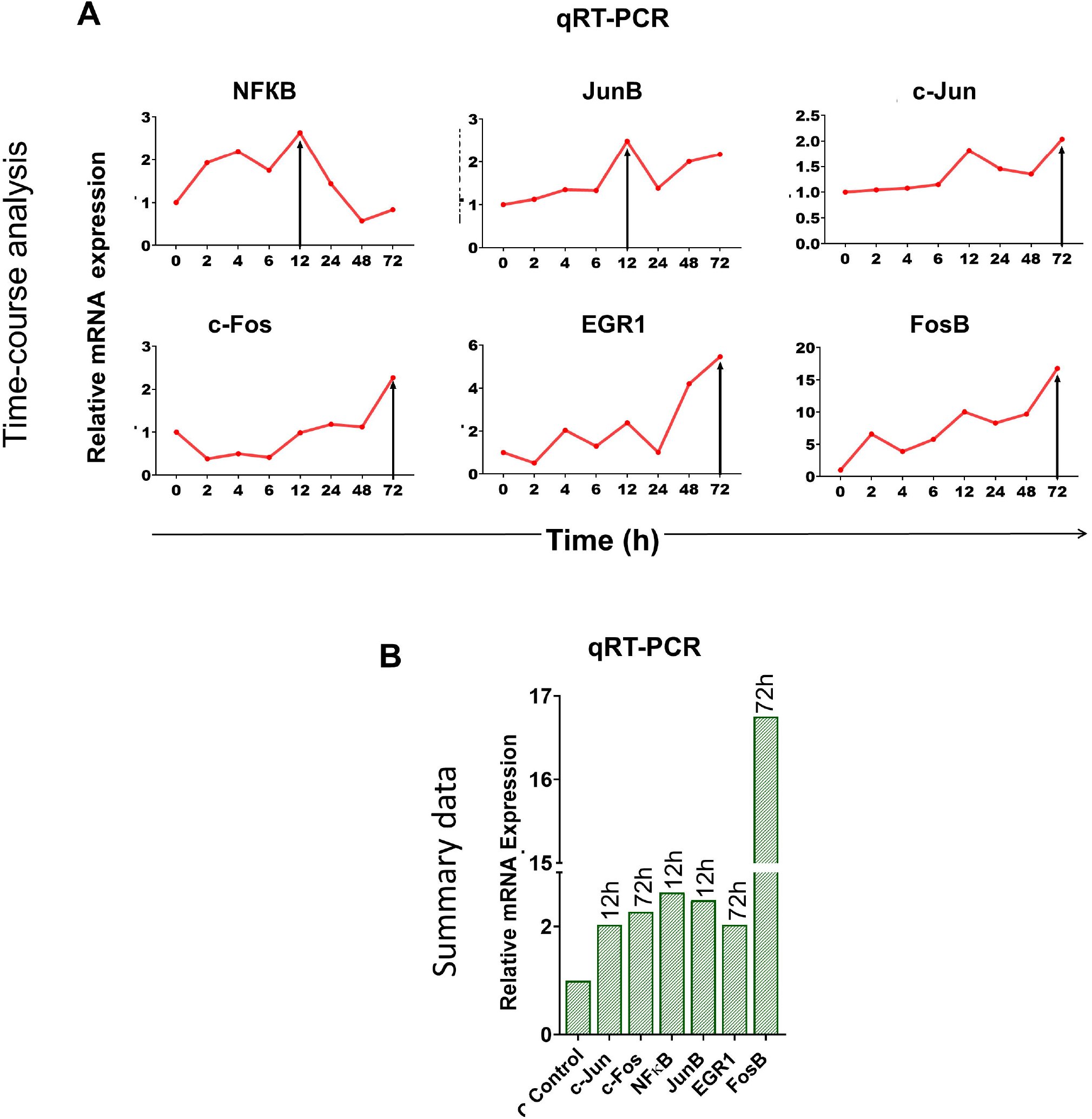
Upregulation of stress-related markers in human PBMCs following treatment of cfChPs isolated from sera of cancer patients. mRNA expression of six stress-related markers in PBMCs was detected by qRT-PCR and analyzed using a comparative C_T_ method. **(A)** Line graphs represent results of time-course analyses calculated as expression fold [2^(-ΔΔC_T_)]. **(B)** Histograms represent relative mRNA expression of stress-related markers at respective peak time points.

### cfChPs released from dying HeLa cells are readily internalized by human lymphocytes

In order to investigate if the above findings could be replicated under a more physiological condition, we undertook the following experiment. HeLa cells that had been fluorescently dual labelled in their DNA and histones were induced to undergo apoptosis in a hypoxia chamber, and the culture medium was passed through a porous membrane (pore size ∼400nm). Human PBMCs were incubated in the filtered culture medium containing dually labelled cfChPs that has been released by the dying HeLa cells (please see Methods). Fluorescence microscopy detected abundant uptake of dually labelled cfChPs by PBMCs and which had accumulated in their nuclei when examined at 4h **(Fig. 4A)**.

**Figure 4:**
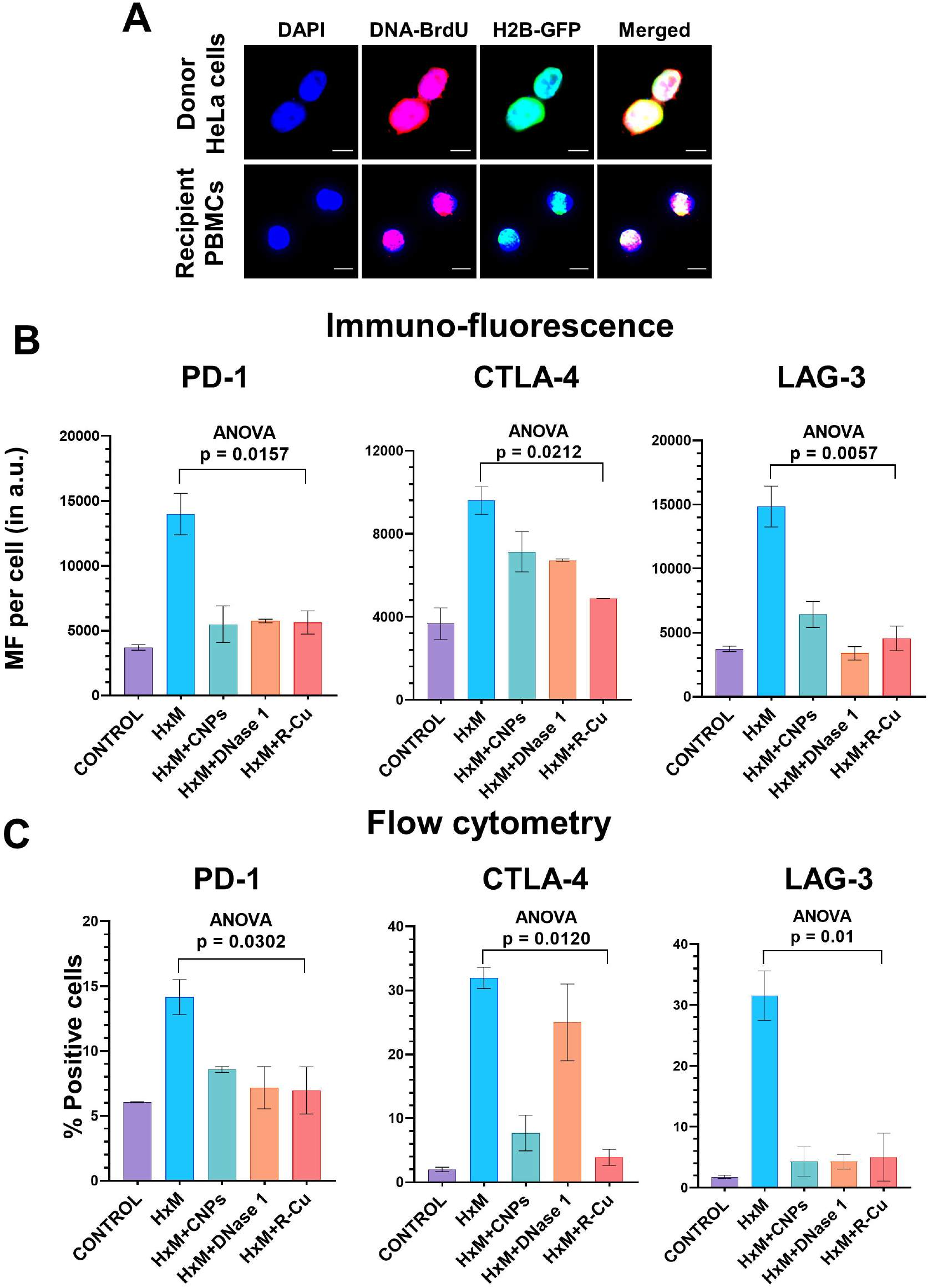
Upregulation of immune checkpoints by cfChPs released from hypoxia-induced dying HeLa cells and their abrogation by cfChPs deactivating agents **(A)** Uptake by PBMCs of fluorescent cfChPs released from fluorescently dually labelled dying HeLa cells. A dual chamber system of Thincert®Cell Inserts was used and cfChPs released from hypoxia-induced dying HeLa cells were collected from the lower chamber (200uL) and added to PBMCs as described in the methods section. **(B)** Results of quantitative IF analysis of upregulation of three immune checkpoints at peak time points of mRNA expression (Figure 3A) and their inhibition by concurrent treatment with the cfChPs deactivators, viz. CNPs, DNase I and R-Cu **(C)** Results of quantitative flow cytometry analysis of upregulation of three immune checkpoints at peak time points of mRNA expression (Figure 3A) and their inhibition by concurrent treatment with the cfChPs deactivators, viz. CNPs, DNase I and R-Cu. Experiments were performed in duplicates. Results are represented as mean ± SEM values and data were analyzed using one-way ANOVA (GraphPad Prism 8).

### Upregulation of immune checkpoints and their abrogated by cfChPs deactivating agents

Having demonstrated that cfChPs from hypoxia-induced dying cells can readily enter into nuclei of human lymphocytes, two hundred and fifty microliters of conditioned culture medium containing hypoxia derived cfChPs were applied to purified T cells. We observed marked up-regulation of all five immune checkpoints and six stress-related markers when examined by qRT-PCR at different time points **(Supplementary Fig. 4)**. In order to confirm that cfChPs in the culture media were the immune checkpoint activating agents, we used the above conditioned media which had been pre-treated with three different cfChPs inactivating agents, namely, anti-histone antibody complexed nanoparticles (CNPs), DNase I and a novel pro-oxidant combination of Resveratrol and copper. Immunofluorescence and flow cytometry analysis were done at the peak time point of mRNA expression as shown in Supplementary Fig. S4 to investigate if the cfChPs inactivating agents would downregulate immune checkpoints that have been approved for clinical use viz. PD-1, CTLA-4 and LAG-3. Highly significant reduction in protein expression with respect to all three immune checkpoints were clearly evident when evaluated by both immunofluorescence and flow cytometry **(Fig. 4B & C; Supplementary Fig. S5 & S6)**.

## Discussion

Following the initial success with immune therapy of cancer, several targeted immune checkpoint inhibitors have been approved for clinical use (4). However, how immune checkpoints are regulated has remained unknown. We have shown here that cfChPs that circulate in the blood of cancer patients, or those that are released naturally from hypoxia-induced dying cancer cells, are the key triggers that activate them. We show that cfChPs are readily internalized by human lymphocytes to rapidly accumulate in their nuclei to activate multiple immune checkpoints in the treated lymphocytes. Our finding of concurrent activation of stress markers leads to the hypothesis that immune activation represents a stress response to DNA damage that results from illegitimate genomic integration of cfChPs (11,13). That activation of immune checkpoints can be abrogated by three different agents that have the ability to deactivate cfChPs, provides further evidence for the involvement of cfChPs in immune checkpoint activation. They also herald the prospect of a novel form of immunotherapy that downregulates multiple immune checkpoints all at once. Such a therapy would be fundamentally different from the current practice of attempting to block already activated immune checkpoints with specific monoclonal antibodies. Above all, cfChPs deactivating agents are likely to be far less toxic and less expensive than the immunotherapeutic agents that are currently in use. Of the three agents, the combination of Resveratrol and copper (R-Cu) is the most promising, since it has already been shown to be effective in multiple therapeutic indications in humans (14–16). No toxic side effects attributable to R-Cu were detected. R-Cu treatment would have the added benefit of reducing toxic side effects in case it was to be used as an adjunct to chemotherapy. Taken together, our results reported in this article, would suggest that R-Cu would be an ideal form of cancer immunotherapy since it can simultaneously down-regulate multiple immune checkpoints with little toxicity and at low cost.

## Methods

### Institutional ethics approval

This study was approved for cfChPs and lymphocyte isolation from blood of healthy volunteers by Institutional Review Board of Advanced Centre for Treatment, Research and Education in Cancer (ACTREC), Tata Memorial Centre (TMC) (Approval no. 900520).

### Animal ethics approval

The experimental protocol of this study was approved by the Institutional Animal Ethics Committee (IAEC) of Advanced Centre for Treatment, Research and Education in Cancer (ACTREC), Tata Memorial Centre (TMC) (Approval no. 12/2020). The experiments were carried out in compliance with the IAEC animal safety guidelines, and with those of ARRIVE guidelines.

ACTREC-IAEC maintains respectful treatment, care and use of animals in scientific research. It aims that the use of animals in research contributes to the advancement of knowledge following the ethical and scientific necessities. All scientists and technicians involved in this study have undergone training in ethical handling and management of animals under supervision of FELASA certified attending veterinarian. Inbred female C57Bl/6 mice were obtained from our Institutional Animal Facility. All mice were maintained in covenant with Institutional Animal Ethics Committee (IAEC) standards. Animals were euthanized at appropriate time points under CO_2_ atmosphere by cervical dislocation under supervision of FELASA trained animal facility personnel.

### Collection of blood samples for cfChPs and lymphocyte isolation

All participants signed a written informed consent form which was approved by the Institutional IRB. For lymphocyte culture, peripheral blood samples from healthy adult volunteers were collected using Vacutainer TM tubes (Becton-Dickinson Vacutainer Systems, Franklin Lakes, NJ, U.S.A.) containing sodium heparin anticoagulant and for cfChPs isolation in plain vacutainer (VACUETTE blood collection tube Serum Clot Activator PREMIUM).

### Isolation of cfChPs from human sera

cfChPs were isolated from sera of cancer patients according to a protocol described by us earlier (11). In order to maintain inter-experimental consistency, pooled serum (typically from ∼5 individuals) was used to isolate cfChPs. cfChPs were quantified in terms of their DNA content as estimated by the Pico-green quantification assay.

### Fluorescent dual labelling of cfChPs

cfChPs were fluorescently dually labelled in their DNA by Platinum Bright 550 (red) and in their histone H4 with ATTO-TEC 488 (green) according to a protocol described by us earlier (11).

### PBMC isolation

Peripheral blood mononuclear cells (PBMC) were isolated using Ficoll-Hypaque according to standard procedures.

### FACs sorting

PBMCs were stained with FITC-conjugated anti-human CD4 antibody (clone PRA-T4, BD Biosciences Pharmingen, San Jose, CA, USA) and PerCP-conjugated anti-human CD8 antibody (clone SK1, BD Biosciences Pharmingen, San Jose, CA, USA) or with FITC-conjugated anti-human CD3 antibody (clone UCHT1, BD Biosciences Pharmingen, San Jose, CA, USA). Cells were sorted on FACS Aria III (BD Biosciences, San Jose, CA, USA). Data were analyzed using FACS Diva software (version 4.0.1.2; Becton, Dickinson and Company). Sorted cells were seeded in 24-well plates and were allowed to rest overnight at 37°C in humidified atmosphere of 5% CO_2_ prior to stimulation.

### Preparation of cfChPs deactivating agents

The cfChPs deactivating agents used in our study were: 1) anti-histone antibody complexed nanoparticles (CNPs) which deactivate cfChPs by binding to histones (17); 2) DNase I from bovine pancreas was procured from Sigma-Aldrich, and 3) a novel pro-oxidant combination of Resveratrol and Copper(R-Cu), which can deactivate / degrade cfChPs via the medium of free-radicals(18). The involved concentrations of R and Cu used in this study were 1mM and 0.0001mM respectively (18). Details of preparation of the cfChPs deactivating agents have been described by us earlier (19).

### Treatment of lymphocytes with cfChPs isolated from sera of cancer patients

Sorted CD4^+^T cells and CD8^+^T cells were plated at a density of 5×10^5^ in 24-well plates containing 1ml of DMEM. After overnight culture, cells were treated with cfChPs (10ng equivalent of DNA) isolated from sera of cancer patients and used for determining upregulation of immune checkpoints.

### Procedure for collecting conditioned media from hypoxia induced dying HeLa cells

A dual chamber system was used to generate conditioned medium containing cfChPs released from dying cells. HeLa cells (∼1×10^5^) were seeded on ThinCert® Cell Culture Inserts (pore size 400nm) containing 1.5ml of DMEM and were placed in 6-well culture plate and were incubated overnight at 37 °C. The six-well plate with Thincert^®^ Inserts was transferred to a hypoxia chamber with 1% O_2_ for 48h to induce hypoxic cell death. Seven hundred microliters DMEM was then added to the lower chamber of the six-well plate below the ThinCert® Inserts and were placed at 37°C in humidified atmosphere of 5% CO_2_ for 48h under normoxic conditions. This procedure allowed cfChPs <400 nm in size released from the hypoxic HeLa cells to seep into the medium in the lower chamber. Post-incubation, conditioned media from the wells was pooled and was used for the co-culture experiments.

### Fluorescent dual labelling of HeLa cells for co-culture experiments

HeLa cells were dually labeled in their histones (H2B) and DNA. Histone H2B labelling was done for 36h with CellLight® Histone 2BGFP (Thermo Fisher Scientific, MA, USA) and DNA labeling was done for 24h using BrdU (10 μM Sigma Chemicals, MO, USA). Dually labeled HeLa cells were seeded on ThinCert® Cell Culture Inserts and cultured in hypoxic conditions (1% O_2_) for 48h. Five hundred micro-liters of conditioned media containing cfChPs <400nm that had seeped into the lower chamber of the insert was applied to isolated PBMCs in a time course experiment (2h, 4h and 6h). Cells were then washed and processed for fluorescence microscopy to detect presence of fluorescent signals of BrdU and histone H2BGFP in the recipient PBMCs.

### Treatment of cells with conditioned media collected from hypoxic HeLa cells

Media from hypoxia treated HeLa cells was collected as described above. Sorted T cells were plated at a density of 5×10^5^ in 24-well plate containing 250μL of DMEM media. After overnight culture, cells were treated with 250μL of conditioned media and a time course analysis using qRT-PCR was performed to determine up-regulation of immune checkpoint. In order to confirm that the active agents in the hypoxic media were cfChPs released from the dying HeLa cells, the hypoxic media was pre-treated for 1h with the following cfChPs inhibitors: 1) CNPs (25µg of anti-H4 IgG conjugated nanoparticles) in 125 µL of phosphate buffer; 2) DNase I in PBS to achieve a final concentration of 0.05U/mL; 3) R:Cu in distilled water to achieve a final molar ratio of 1 mM R : 0.0001 mM Cu).

### Analyses of immune checkpoints

#### qRT-PCR

Evaluation of immune checkpoints was performed using qRT-PCR following treatment of human lymphocytes with cfChPs or with media of hypoxic dying HeLa cells as described above. The time points for analyses were: 0h, 30min, 6h, 12h, 24h, 36h and 48h. Appropriate untreated control cells for each time point were analysed in parallel. Total RNA was isolated using RNeasy Mini Kit (Qiagen, Hilden, Germany) and approximately 1 µg of isolated RNA was converted to cDNA using RT^2^ First Strand Kit (Qiagen, Hilden, Germany). cDNA was diluted (1:10) and used in 10µl reaction volume in duplicates. Real-time PCR was carried out using SYBR Select Master Mix (Applied Biosystems, CA, USA) and all the samples were assayed on a QuantStudioTM 12K Flex Real-Time PCR System (ThermoFisher) using a 384-well block in duplicates. Data were analysed using a comparative C_T_ method and fold change in mRNA expression was calculated as 2^(-ΔΔC_T_).

#### Immunofluorescence

For detection of immune checkpoints expression, immunofluorescence analysis for five immune checkpoints was performed at peak time points of mRNA expression following treatment of cells. Methodological details of immunofluorescence have been described by us earlier (10). Briefly, slides were mounted with vectashield mounting medium with DAPI (Vector Laboratories) and analyzed on Applied Spectral Imaging system as described by us earlier (10). All experiments were performed in duplicate, and 500 cells were analysed in each case. Results were expressed as mean ± SEM.

#### Flow cytometry

After cfChPs treatment, PBMCs were labelled with the following antibodies; anti-CD3-FITC (clone:UCHT1), anti-CD-8 PerCP (clone:SK1), anti-PD-1-BV421 (clone:EH12.1), anti-CTLA-4-PE-CF594 (clone:BN13), anti-LAG-3-BUV395 (clone:T47-530), anti-NKG2A-BV421 (clone:131411) and anti-TIM-3-BV786 (clone:7D3). All antibodies were purchased from Becton Dickinson Biosciences, San Jose, CA, USA. Cells were incubated for 20mins in dark at room temperature followed by PBS wash. Samples were acquired on FACSAria III cytometer (Becton Dickinson, USA) and analysed with *FlowJo*™ *v10*.*6 Software* (ThreeStar Inc, USA). Lymphocytes were gated on forward and side scatter parameters. At least 20,000 cells were acquired and the threshold between negative and positive expression was defined by the fluorescence minus one (FMO) method to determine percent positive cells.

### *In vivo* studies

#### Intravenous injection of cfChPs into mice

For the *in vivo* study, 21 mice (3 per group, 7 groups) were injected with cfChPs (100ng dissolved in saline for each mouse) and 3 mice acted as untreated controls. Mice were sacrificed under anesthesia at 6h, 12h, 18h, 24h, 48h, 72h and 96h after cfChPs injection and spleen was removed.

#### Preparation of mouse splenocytes

The spleens were minced with a sharp sterile blade, placed in a 40-m nylon cell strainer and pressed with the plunger of a syringe. Splenocytes were suspended in RPMI-1640 supplemented with 5% FBS. Red blood cells were lysed with 1XBD Pharmlyse, washed, and splenocytes were re-suspended in 5% FBS in PBS.

#### Analyses of activation of immune checkpoints in splenocytes by flow cytometry

One million splenocytes were stained for 20 min in the dark with the following antibodies: CD3-APC-Cy7 (Clone: 17A2), CD4-FITC (Clone:GK1.5), CD8-APC (Clone:53-6.7), PD-1-BV510 (Clone:29F.1A12), CTLA-4-PE (Clone:UC10-4B9), NKG2A-PE (Clone:16A11), Tim-3-BV421 (Clone:RMT3-23) and LAG-3-PerCP-Cy5.5 (Clone:C9B7W). All antibodies were purchased from BioLegend company (USA). Samples were acquired on BD FACS Aria III (Becton Dickinson, USA) and analysed with *FlowJo*™ *v10*.*6 Software* (ThreeStar Inc, USA) as described above.

### Statistical analysis

All data are presented as Mean□±□Standard Error of Mean (SEM). Statistical analysis was performed using GraphPad Prism 8 (GraphPad Software, Inc., USA, Version 8.0). Data were compared using Student’s□t-□test (two tailed, unpaired), one-way ANNOVA and Bonferroni’s multiple comparisons test. *p*□<□0.05 was taken as the level of significance.

## Acknowledgments

The authors thank flow cytometry facility at ACTREC-TMC for their technical support. The authors thank Dr. Rajan Basak for his guidence in qRT-PCR.

## Authors contributions

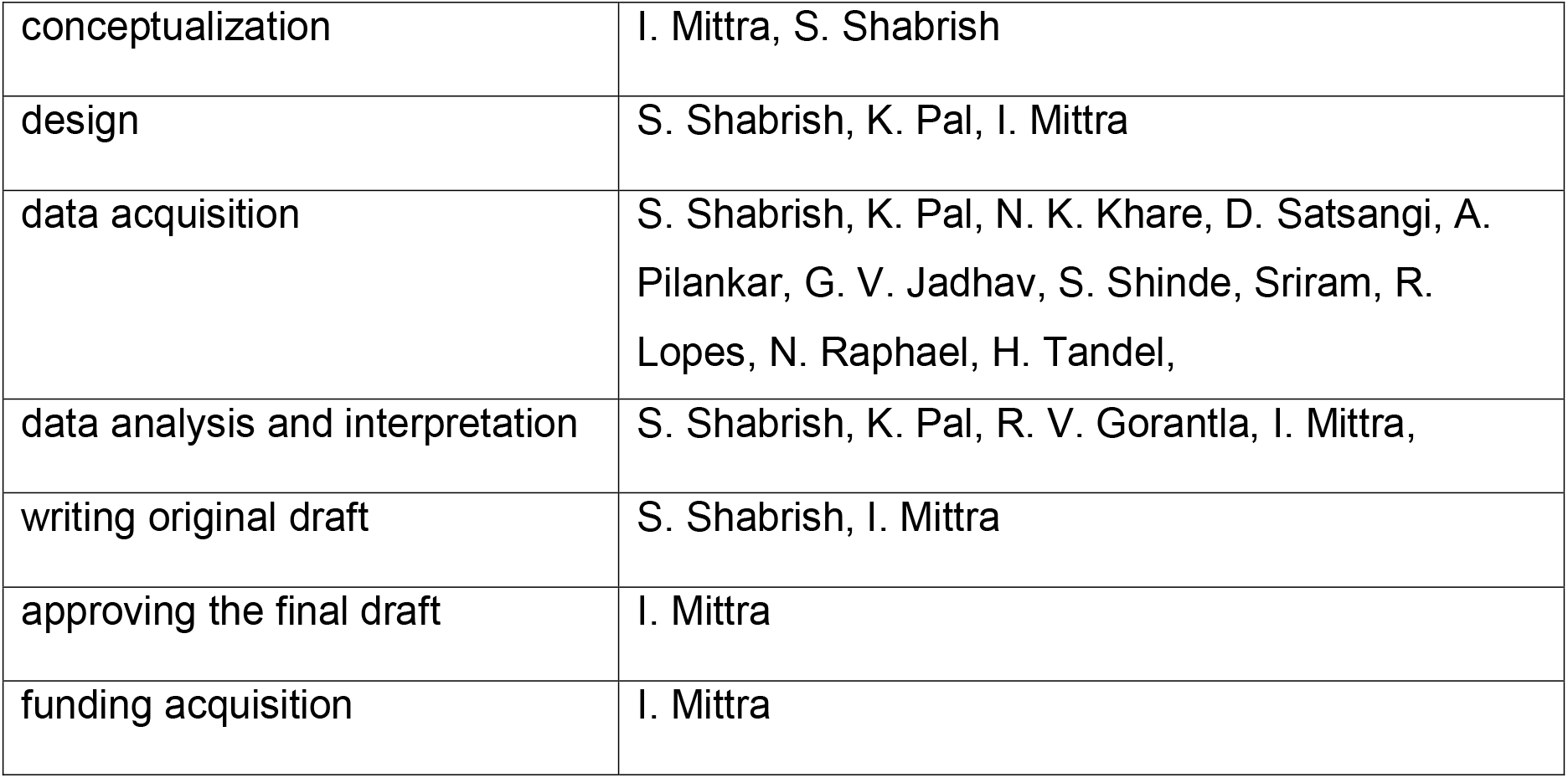

## Data and materials availability

All data are available in the manuscript and in supplementary material.

## Legends to supplementary figure

**Supplementary Fig. 1:** Rapid internalization by human PBMCs of cfChPs isolated from sera of cancer patients. Fluorescently dually labeled cfChPs (10ng) when added to PBMCs are seen to have accumulated in their nuclei at 2h.

**Supplementary Fig. 2:** Representative IF images showing upregulation of five immune checkpoints in **(A)** CD4^+^T cells and **(B)** CD8^+^T cells following treatment with cfChPs isolated from sera of cancer patients. Scale bars = 20µm.

**Supplementary Fig. 3:** Representative flow cytometry plots showing upregulation of surface expression of immune checkpoint expressions on **(A)** CD4^+^T cells and **(B)** CD8^+^T cells following treatment with cfChPs isolated from sera of cancer patients.

**Supplementary Fig. 4:** Time-course analysis using qRT-PCR to determine the point of maximum expression of immune checkpoints and stress markers following treatment of PBMCs with conditioned media containing cfChPs released from hypoxia-induced dying HeLa cells. **(A)** Upregulation of immune checkpoints and **(B)** upregulation stress related markers.

**Supplementary Fig. 5:** Representative IF images showing immune checkpoint expression in T cells following treatment of PBMCs with conditioned media containing cfChPs released from hypoxia-induced dying HeLa cells. Pre-treatment of conditioned medium with CNPs (25 ug), DNase I (0.005U) and R-Cu (1:10^−4^) significantly reduced the expression of immune checkpoints.

**Supplementary Fig. 6:** Representative flow cytometry plots showing immune checkpoint expression in T cells following treatment of PBMCs with conditioned media containing cfChPs released from hypoxia-induced dying HeLa cells. Pre-treatment of conditioned medium with CNPs (25 μg), DNase I (0.005U) and R-Cu (1:10^−4^) significantly reduced the expression of immune checkpoints.

