## Supplementary figures and images for "Cell-free chromatin particles activate immune checkpoints in human T cells: Implications for cancer therapy"

### Supplementary Figure 1

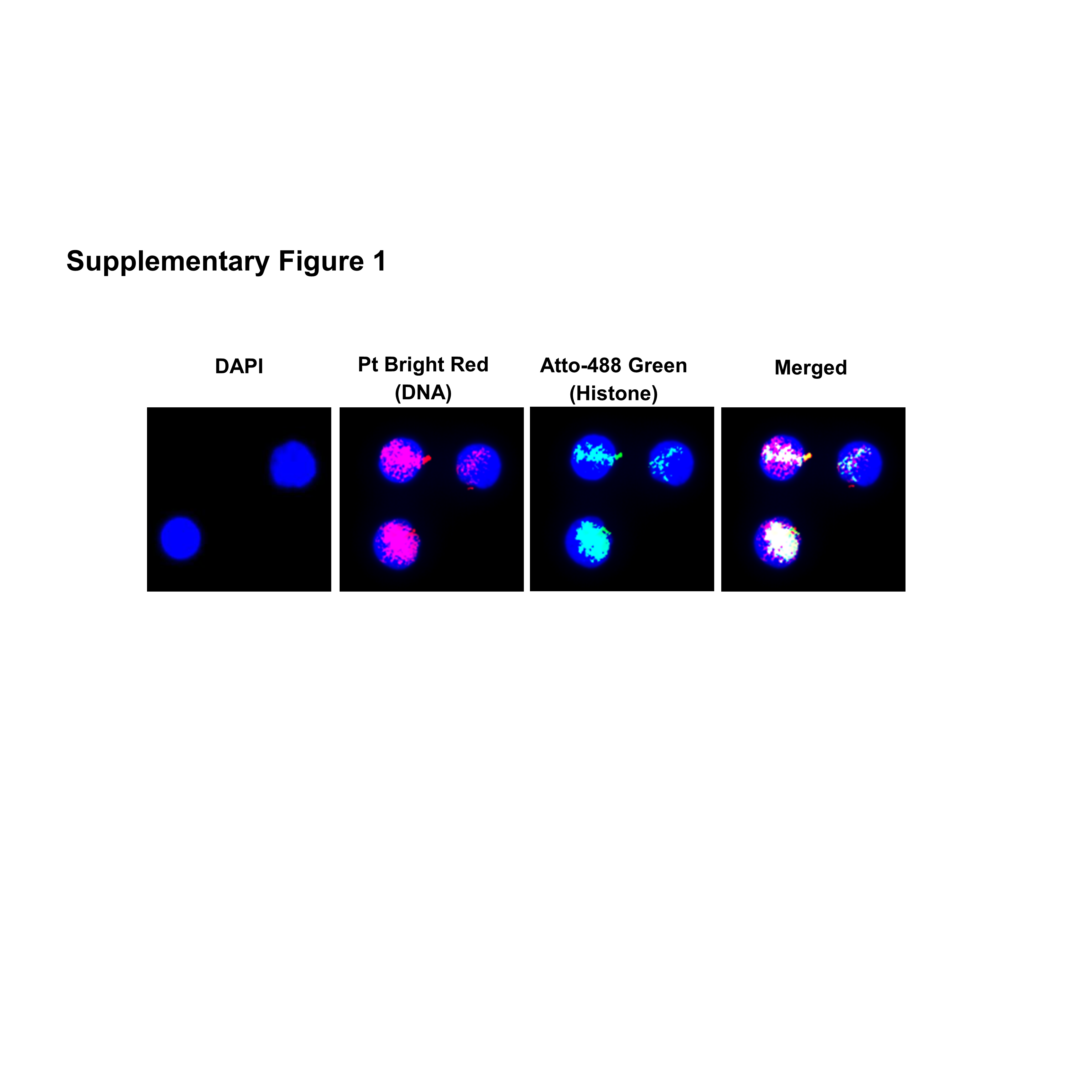

### Supplementary Figure 2

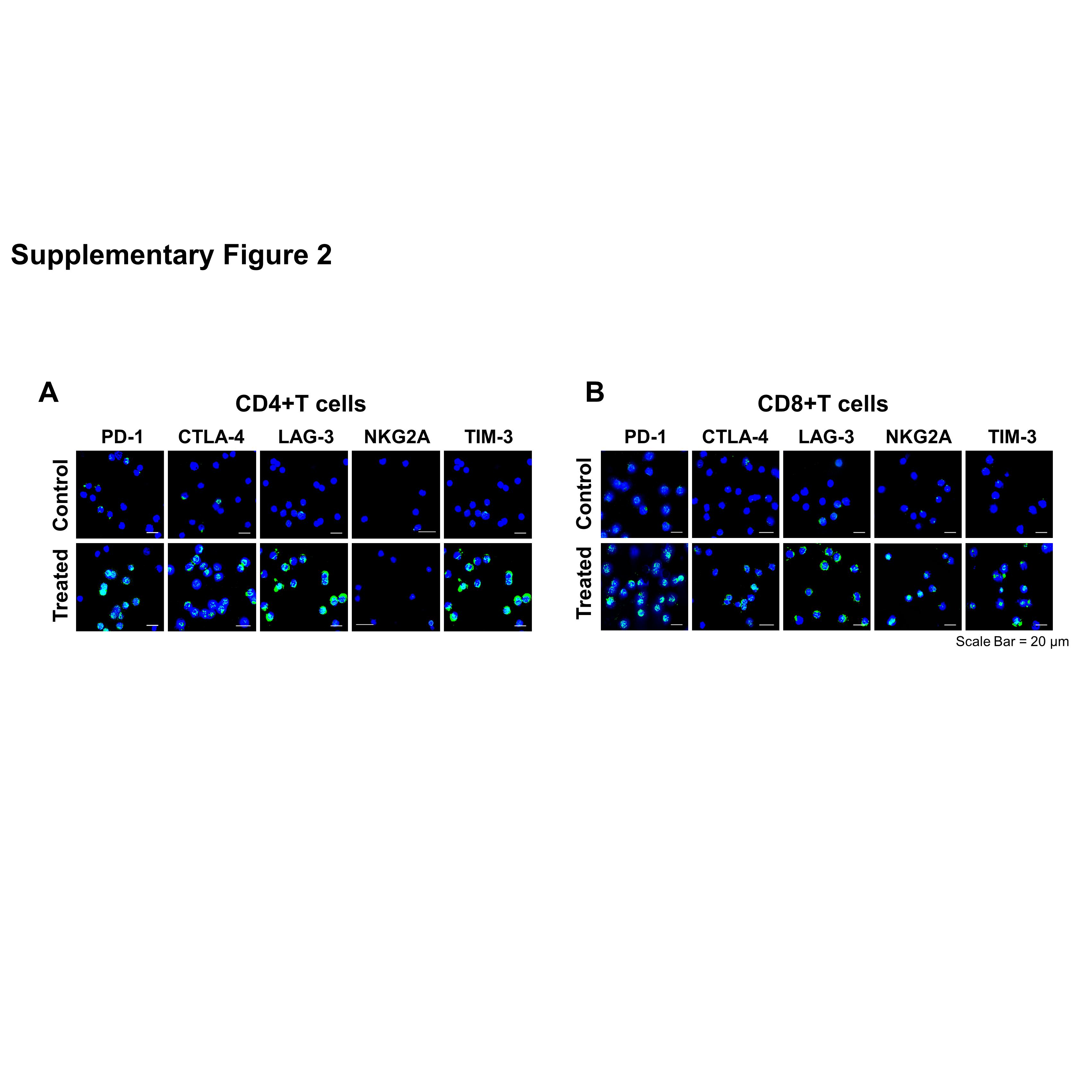

### Supplementary Figure 3

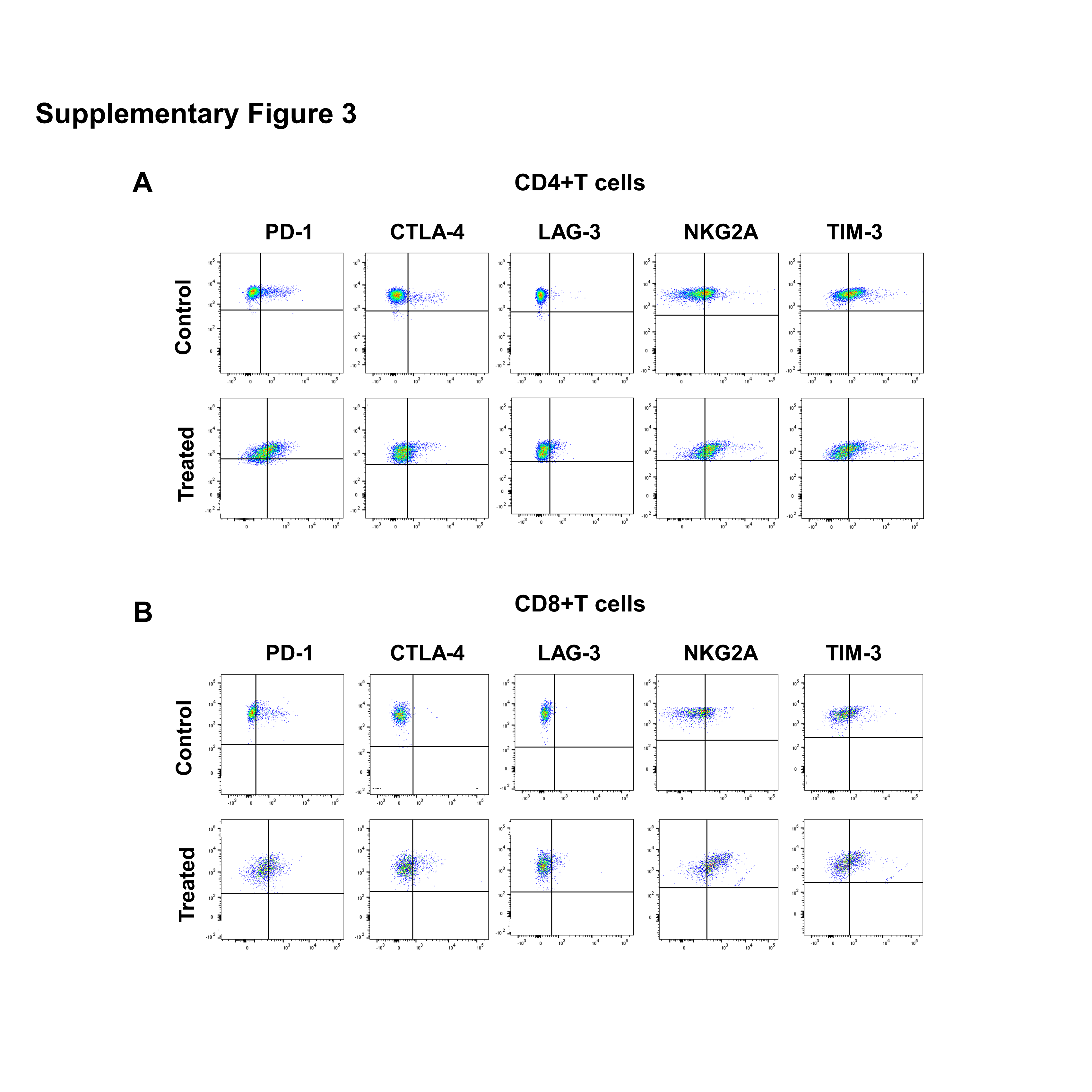

### Supplementary Figure 4

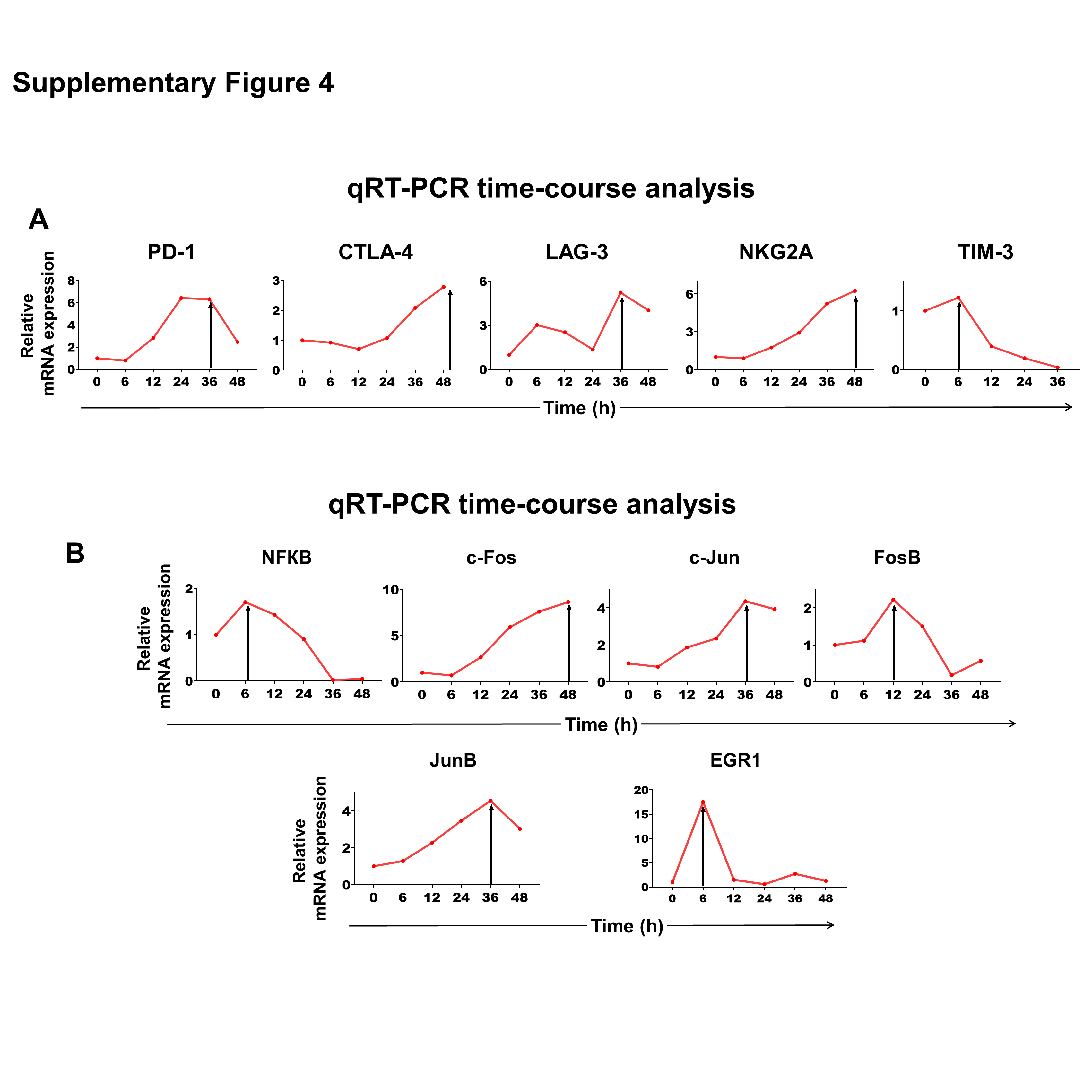

### Supplementary Figure 5

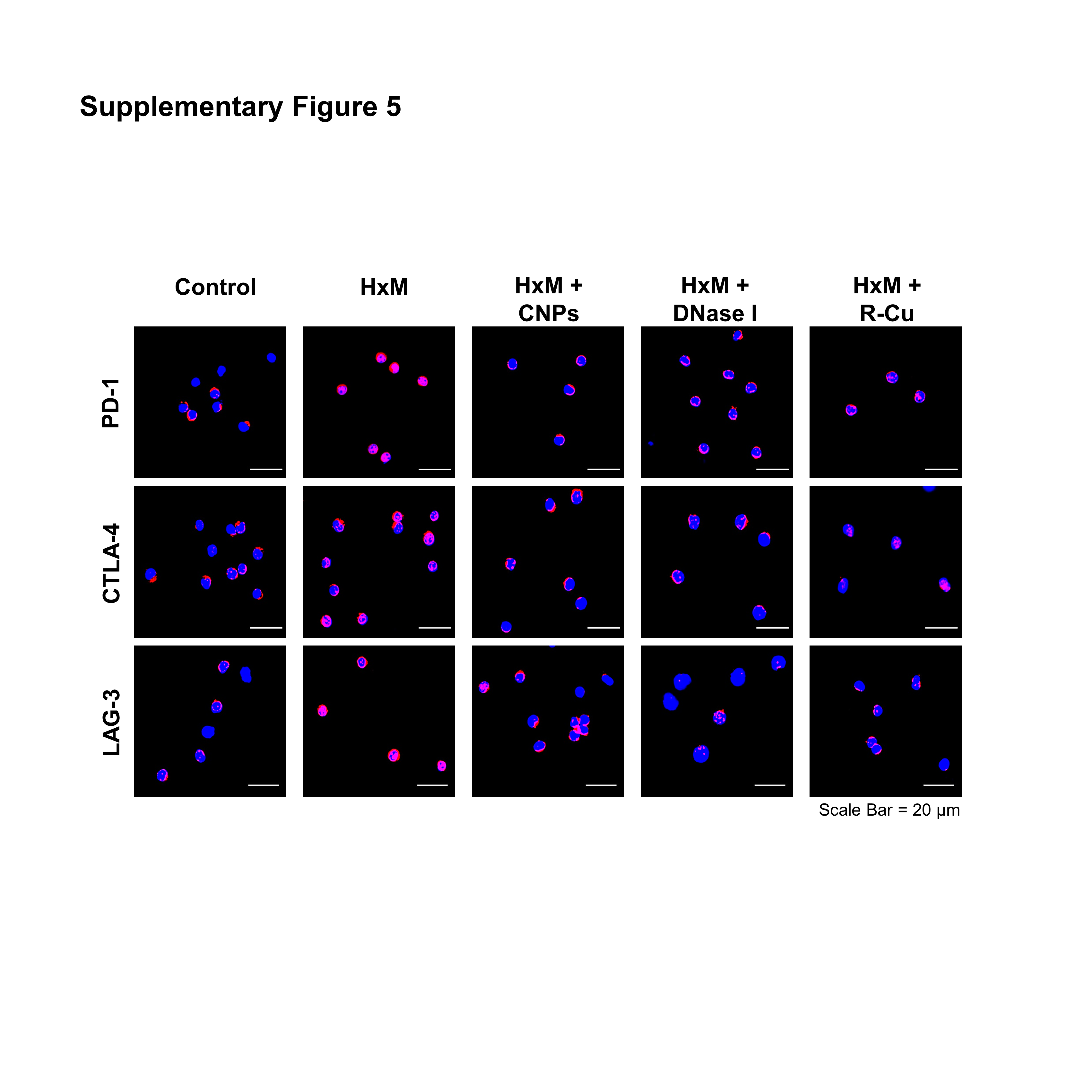

### Supplementary Figure 6

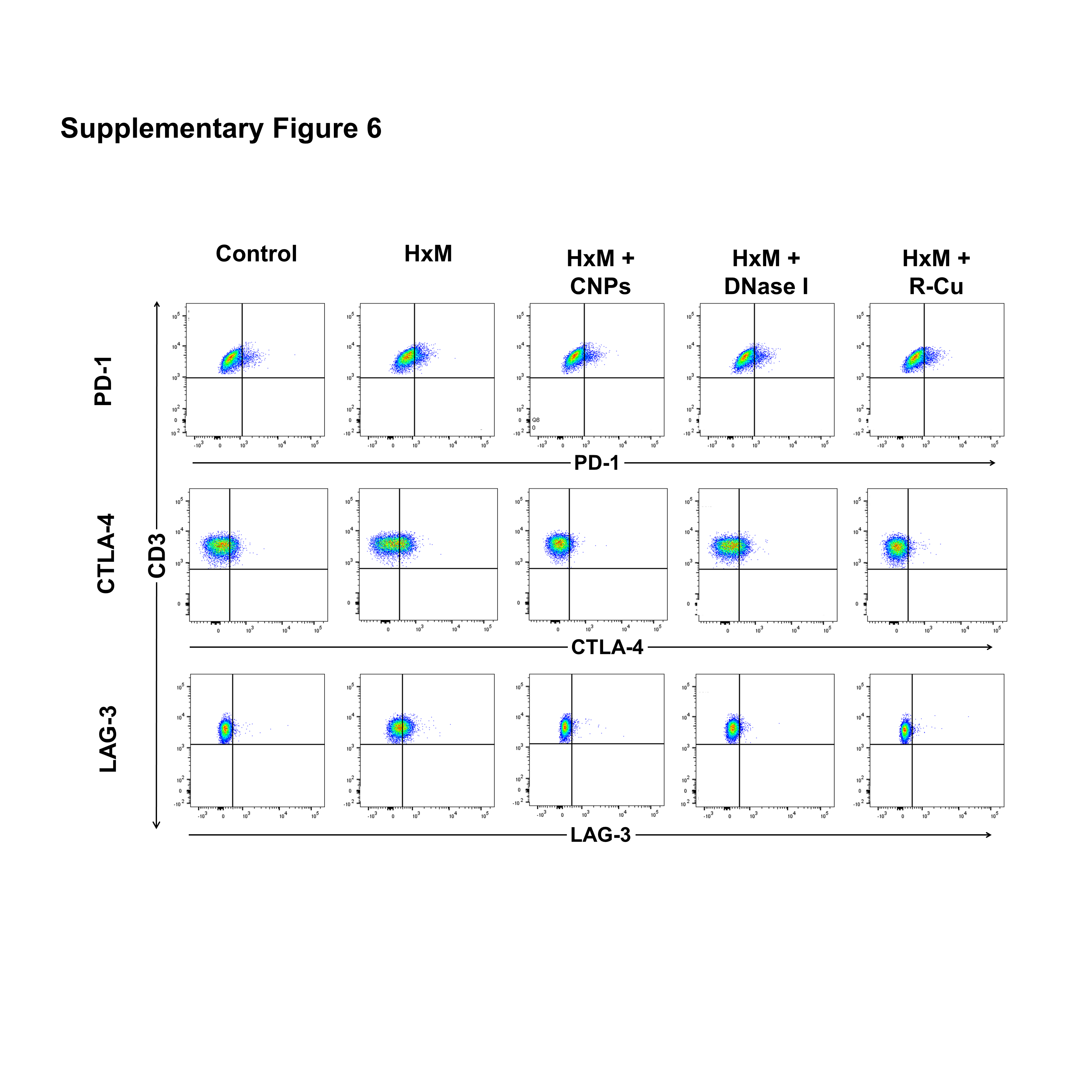
